# Effect of ceftiofur cessation and substitution with lincomycin-spectinomycin on extended-spectrum-beta-lactamase/AmpC genes and multidrug resistance in *E. coli* from a Canadian broiler production pyramid

**DOI:** 10.1101/521120

**Authors:** L. Verrette, J. M. Fairbrother, M. Boulianne

## Abstract

Ceftiofur, a cephalosporin antimicrobial, was used systematically in Canadian hatcheries for many years to prevent early mortality in chicks leading to a high prevalence of cephalosporin resistance in *Escherichia coli* in chickens. Preventive use of ceftiofur in hatcheries ceased in 2014. We examined the effect of ceftiofur cessation and replacement with lincomycin-spectinomycin at the hatchery on the proportion of *E. coli* positive for extended-spectrum β-lactamase (ESBL) and AmpC β-lactamase related genes, and on the multidrug resistance profiles of ESBL/AmpC positive *E. coli* in broilers and their associated breeders, at one year post-cessation. For indicator *E. coli* from non-enriched media, a significant decrease post-cessation in the proportion of samples harboring *E. coli* isolates positive for *bla*_CMY-2_ and/or *bla*_CTX-M_ was observed. In contrast, following enrichment in medium containing ceftriaxone (1mg/L) to facilitate recovery of ESBL/AmpC β-lactamase producing *E. coli* colonies, both pre- and post-cessation, 99% of the samples harbored *E. coli* positive for *bla*_CMY-2_ or *bla*_CTX-M_. Flocks receiving lincomycin-spectinomycin after cessation of ceftiofur showed a significantly greater non-susceptibility to aminoglycoside, folate inhibitor, phenicol, tetracycline and possible extensively drug resistant *E. coli* compared to those receiving ceftiofur or no antimicrobial at hatchery. This study clearly demonstrates an initial decrease in ESBL/AmpC positive *E. coli* following the cessation of ceftiofur in hatchery but an increase in multidrug resistant *E. coli* following replacement with lincomycin-spectinomycin.

**Importance:** Antimicrobial resistance is a global problem. The antimicrobial ceftiofur has been used worldwide for disease prevention in poultry production resulting in a greatly increased resistance to this antimicrobial important in poultry and human medicine. Our study examines the impact of ceftiofur cessation and its replacement with the antimicrobial combination lincomycin-spectinomycin, a common practice in the industry. Our study demonstrated a decrease in ceftiofur resistance after the cessation of its use, although the resistance genes remain ubiquitous in all phases of poultry production, showing that poultry remains a reservoir for ceftiofur resistance and requiring continued vigilance. We also observed a decrease in multidrug resistance after cessation of ceftiofur although the contrary finding following use of lincomycin-spectinomycin indicates that the use of these antimicrobials should be questioned. Reduced resistance to ceftiofur in poultry may translate to better treatment efficacy, decreased morbidity, mortality, duration and cost of hospitalization in humans.

## Introduction

One of the most important causes of early mortality in broiler chick is omphalitis, mostly caused by *avian pathogenic E. coli* (APEC), a subgroup of *extra-intestinal pathogenic E. coli* (ExPEC) (1, 2). Ceftiofur, a third generation cephalosporin antimicrobial, has been administered for over 15 years either *in ovo* or by subcutaneous injection at the hatchery, in order to reduce early chick mortality in many countries (3) Consequently, an increased prevalence of extended-spectrum β-lactamase (ESBL) and AmpC β-lactamase producing *Escherichia coli* has been reported worldwide (4-7), resulting in an increased extended spectrum cephalosporin resistance in the broiler poultry chain production. This is a public health concern due to co-resistance with other extended spectrum cephalosporins, such as ceftriaxone and cephamycin, antimicrobials which are used widely in human medicine (8, 9). ESBL/AmpC-associated resistance genes detected in chickens are *bla*_CMY-2_, *bla*_SHV,_ *bla*_CTX-M,_ *bla*_OXA,_ and *bla*_TEM_ (8, 10). Ceftiofur has been used in food-producing animals in North America since 1989, and *bla*_CMY-2_ gene was first reported in 1998 from cattle (11).

In Canada in 2014, the poultry industry eliminated the preventive use of Ceftiofur in hatcheries for the second time (3). Following the first cessation in 2005, a decline in the prevalence of cephalosporin-resistant *Salmonella Heidelberg* isolates in chicken meat was observed, although the effect on prevalence of resistance in *E. coli* was not clear (9). Recent studies have shown a decrease in the proportion of clinical isolates possessing ESBL/AmpC-associated resistance genes after this second cessation (6, 12, 13). In addition, the prevalence of resistant *E. coli* from healthy broilers at farms was markedly decreased within a year after ceftiofur cessation at hatcheries in Japan (4). A decrease in the prevalence of *Salmonella* harboring *bla*_CMY-2_ in chicken meat was also observed in Japan in the same years (14). In Canada, whereas some hatcheries completely stopped using antimicrobials following the cessation of ceftiofur (15), other hatcheries replaced it with lincomycin-spectinomycin (3, 6).

To our knowledge, there has been no convenient sampling study of healthy flocks in Canada comparing the impact of ceasing the administration of ceftiofur or any other antimicrobial *in ovo* and replacing it with lincomycin-spectinomycin at the hatchery. Hence, our objectives were to examine the effect of ceftiofur cessation and replacement with lincomycin-spectinomycin at the hatchery on the proportion of *E. coli* possessing the ESBL/AmpC genes *bla*_CMY-2_, *bla*_SHV,_ *bla*_OXA,_ *bla*_CTX-M_ and *bla*_TEM_, and on the multidrug resistance profiles of these isolates in young chicks, broilers and breeders of an integrated pyramid.

## Material and Methods

### 1. Sampling

Two vertical samplings of meat chickens of an integrated pyramid in the province of Quebec in Canada were made. The first sampling period was between March and May 2014 when chicks were routinely receiving 0,08-0,2 mg of ceftiofur per egg as an *in ovo* injection (16, 17). The second sampling was done between June and October 2015, one year after preventive administration of ceftiofur at the hatchery had ceased. Chicks on about half of the farms in the 2015 sampling period received lincomycin and spectinomycin (2,5 mg of lincomycin and 5 mg of spectinomycin per chick (17)) *in ovo* whereas chicks on the remaining farms did not receive any preventive antimicrobial at the hatchery.

#### a. Breeding sampling

Eight broiler breeder flocks belonging to the same hatchery were conveniently selected for both samplings. For both sampling years, three successive samplings were made in the breeder flocks, within one month (+/-one week). For fecal sampling, the floor of each breeder house was divided into four quarters, and approximately ten fresh fecal droppings were collected from each quarter, put on ice and delivered to the EcL laboratory of the Faculty of Veterinary Medicine at Saint-Hyacinthe, where they were kept overnight at 4 °C. The next day, feces of each quarter were mixed manually and 10 g of feces were mixed in 45 ml of peptone water (Oxoid Canada, Nepean, Ontario, Canada). After standing for 30 minutes, 8,5 ml of the peptone water suspension was collected and frozen with 1,5 ml of glycerol at −80 °C.

#### b. Broiler sampling

In the month following the beginning of the sampling of the eight broiler breeders, chicks from corresponding sampled breeder flocks were selected for sampling at hatch and at the end of the growing period. A list of farms was obtained from the hatchery company. The first farmers who agreed to participate were recruited. Study farms housed 5000 to 30 000 chickens. All eggs were incubated at the same hatchery. Shortly after hatch and placement into the delivery box, the paper under the chicks was collected to sample meconium. Papers were delivered to the EcL laboratory, where they were kept overnight at 4 °C. The next day, 8 to 10 pieces of 3 cm by 3 cm cardboard containing meconium were cut out and put in 30 ml of peptone water. The mixture was then incubated at 37 °C overnight. On the next day, 8,5 ml of the peptone water solution was mixed with 1,5 ml of glycerol and frozen at −80 ° C. Upon chick delivery at the farm, care was taken to place the sampled chicks in a single and well identified pen to ensure traceability. Between 18 and 29 days of age, fecal sampling was done for each flock using the protocol previously described for the broiler breeder fecal sampling.

In 2014, during ceftiofur use, a total of 22 composite fecal samples from the 8 breeder flocks (one flock was sampled only once before going to slaughter), 20 composite meconium samples (4 meconium samples each allowed the production of two different broiler flocks, making a total of 24 broiler flocks with 20 meconium samples) and 20 composite fecal samples of broiler flocks (traceability to the breeders was lost for 4 lots) were selected. In 2015, a total of 24 samples from 8 breeder flocks, with corresponding meconium and composite fecal samples from 14 broiler chicken flocks for which no *in ovo* antimicrobials were administered and 16 flocks for which lincomycin-spectinomycin was administered *in ovo*, were taken.

### 2. Colony isolation of *Escherichia coli*

From the frozen samples, two different isolation protocols were used to obtain an indicator *E. coli* isolate collection and a potential ESBL/AmpC-producing isolate collection.

#### a. Indicator *E. coli* isolate collection

Samples were spread with a swab on MacConkey agar (Becton Dickinson and Company). After overnight incubation at 37 °C, five well-isolated lactose-positive colonies, when possible, were inoculated on MacConkey and incubated at 37 °C overnight. Five well-isolated colonies were then selected and incubated in Luria–Bertani (LB) (Becton Dickinson and Company) broth overnight at 37 °C. Finally, 750 μL of the bacterial suspension for each isolate was mixed with 750 μL of 30% glycerol and frozen at −80 ° C.

#### b. Potential ESBL/AmpC-producing isolate collection

We used the protocol described previously by Agersø and colleagues with some modifications (18, 19). A volume of 50 μl of each thawed sample was inoculated in 5 ml of peptone water containing 1 mg/ml ceftriaxone (20). After incubation at 37 °C overnight, cultures were streaked on MacConkey agar plates with 1 mg/ml ceftriaxone. After another overnight incubation at 37 °C, 5 colonies were selected (preferably lactose +) and inoculated on MacConkey agar containing 1 mg/ml ceftriaxone overnight. Five well isolated colonies on agar were placed in LB culture medium and incubated overnight at 37 °C. Finally, 750 μL of each bacterial suspension was mixed with 750 μL of 30% glycerol and frozen at −80 °C.

### 3. DNA extraction and *uidA* PCR

For each isolate, DNA was extracted from the overnight culture in LB medium. DNA templates were prepared from the samples by boiled cell lysis for examination by PCR, as described previously by Maluta et al. (21). All isolates were confirmed as *E. coli* by PCR for detection of housekeeping gene *uidA*, which encodes Beta-glucuronidase, with control strain ECL7805 (22). PCR conditions used to detect *uidA* gene included initial denaturation (95°C, 2 min), 24 cycles of denaturation (94°C, 30 sec), annealing (65°C, 30 sec), extension (72°C, 30 sec), and final extension (4°C).

### 4. Detection of antimicrobial resistance genes

*Escherichia coli* isolates positive for *uidA* were analyzed for the presence of *bla* genes by multiplex PCR. Five β-lactamase resistance genes (*bla*_SHV_, *bla*_TEM_, *bla*_CMY-2_, *bla*_OXA_, *bla*_CTX-M_) were tested in a subset of *E. coli* isolates from the potential ESBL/AmpC-producing isolate collection (3 of 5 isolates maximum per sample for a total of 432 tested) and for all the *E. coli* isolates in the indicator collection (5 colonies maximum per sample with a total of 722 tested). The protocol was provided by the National Microbiology Laboratory of the Public Health Agency of Canada and used with some minor adjustments and with the control strains ECL3482, PMON38 and CTX-M15 (23).

### 5. Phenotypic antimicrobial susceptibility testing in the potential ESBL/AmpC-producing isolate collection

For each sample from the potential ESBL/AmpC-producing isolate collection, the first 2 of 3 isolates tested by PCR were selected for examination by the disk diffusion (Kirby Bauer) assay (total of 290 isolates), as previously described by the Clinical and Laboratory Standards Institute (CLSI). Susceptibility of isolates was tested for 14 antimicrobials belonging to 10 classes as used in the Canadian Integrated Program for Antimicrobial Resistance Surveillance (CIPARS) for food producing animals and agents of interest in human and veterinarian medicine (24), with addition of spectinomycin (total of 15 antimicrobials and 10 classes). Breakpoints were those recommended by CLSI in 2015 for Enterobacteriaceae, for most of the antimicrobials (25). There were three exceptions where recommendations in the CLSI in 2015 for animals were used: tetracyclines and ceftiofur, where the breakpoints selected were for Enterobacteriaceae, and spectinomycin, where the zone diameter interpretative standards used were for *Pasteurella multocida* (26). Intermediate and resistant strains together were classified as non-susceptible. *Escherichia coli* strain ATCC 25922 was used as quality control for susceptibility testing.

Multi-drug resistance (MDR) was defined as non-susceptibility to at least one agent in three different antimicrobial classes and possible extreme drug resistance (XDR) as non-susceptibility to at least one agent in all but two or fewer antimicrobial classes tested, as suggested by Magiorakos et al. (27). In order to more precisely compare the level of MDR between groups, the level of multi-drug resistance for each sample was classified from 0 to 10, representing the number of antimicrobial classes to which the sample was non-susceptible.

### 6. Statistical Analysis

The unit of interest for statistical analysis was the flock, one pooled sample representing one flock. A sample was considered non-susceptible to an antimicrobial when at least one of its isolates demonstrated non-susceptibility. The associations between the various groups (*in ovo* administration of Ceftiofur, no antimicrobial or lincomycin-spectinomycin) were tested using exact chi-square with SAS v.9.3 (Cary, N.C.). The alpha value was set at 0,05.

## Results

### 1. Decreased proportion of non-enriched samples with *E. coli* positive for *bla*_CMY-2_ or *bla*_CTX-M_ following cessation of *in ovo* administration of ceftiofur in hatchery

Before the cessation of Ceftiofur, the proportion of samples with *E. coli* positive for *bla*_CMY-2_ was very high for the meconium (90%), decreasing to 60% in the feces of broilers at the end of fattening, and to 0% in breeders (Table 1). Similarly, the proportion of samples with *E.coli* positive for *bla*_CTX-M_ was higher for the meconium (20%) than in broilers (5%) and in breeders (0%). Interestingly, following cessation of ceftiofur and without the *in ovo* administration of lincomycin-spectinomycin, a decrease in the proportion of samples with *E. coli* positive for *bla*_CMY-2_ was observed in the meconium and broiler feces, being significant (p=0,002) only for the meconium. The same trend was observed when comparing the proportion of samples with *E. coli* positive for *bla*_CTX-M_ in meconium and broiler feces, although this result is not statistically different. In 2015, the breeders (which produce the flocks receiving no antimicrobial or lincomycin-spectinomycin *in ovo*) had a low proportion of positive samples to *bla*_CMY-2_ (16%) and *bla*_CTX-M_ (0%). The proportions of meconium and broiler samples in 2015 with *E. coli* positive for *bla*_CMY-2_ or *bla*_CTX-M_ were not significantly different when comparing flocks receiving lincomycin-spectinomycin to those which received no antimicrobial *in ovo*. Following the cessation of ceftiofur, as observed in flocks not receiving any antimicrobial *in ovo*, birds receiving lincomycin-spectinomycin showed a significant decrease in the proportion of samples harboring *E. coli* possessing the *bla*_CMY-2_ gene (p<0,01) for the meconium and a decrease, although not significant, in the proportion of samples harboring *E. coli* possessing *bla*_CMY-2_ for the broiler feces. No significant differences were observed between groups for the proportion of samples with *E. coli* positive for the *bla*_TEM_ gene. Overall, the proportion of samples with *E. coli* positive for the *bla*_SHV_ gene was very low and the results are not shown. No samples with *E. coli* positive for the *bla*_OXA_ gene were detected.

**Table 1:** Effect of cessation of *in ovo* administration of ceftiofur and replacement with lincomycin-spectinomycin on the proportion of non-enriched samples from newly hatched, broiler and breeder birds with *E. coli* isolates positive for ESBL/AmpC resistance genes

| Sample origin | Before or after<br>ceftiofur cessation<br>in broiler | *In ovo*<br>administration<br>of: | No of<br>samples<br>(isolates)<sup>a</sup> | % samples with one or more<br>isolates positive for: |  |  |
| --- | --- | --- | --- | --- | --- | --- |
|  |  |  |  | *bla*<sub>TEM</sub> | *bla*<sub>CMY-2</sub> | *bla*<sub>CTX-M</sub> |
| Meconium of<br>newly hatched<br>chicks | Before (2014) | Ceftiofur | 20 (96) | 45 | 90<sup>b</sup> | 20 |
|  | After (2015) | Nothing | 14 (69) | 71 | 36<sup>c</sup> | 0 |
|  | After (2015) | Lincomycin<br>Spectinomycin | 16 (80) | 56 | 50<sup>c</sup> | 0 |
| Pooled feces of<br>broilers | Before (2014) | Ceftiofur | 20 (100) | 50 | 60 | 5 |
|  | After (2015) | Nothing | 14 (70) | 43 | 50 | 0 |
|  | After (2015) | Lincomycin<br>Spectinomycin | 16 (79) | 69 | 44 | 6 |
| Pooled feces of<br>breeders | Before (2014) | Ceftiofur | 22 (109) | 64 | 0 | 0 |
|  | Before (2015) | Ceftiofur | 24 (119) | 60 | 16 | 0 |

<sup>a</sup> All isolates tested were uidA positive (likely *E. coli*)
<sup>bc</sup> Different letters in superscript in the same column and same sample indicate significantly different results

### 2. High proportion of ceftriaxone-enriched samples from all sources before and after cessation of *in ovo* administration of ceftiofur harbor *E. coli* positive for *bla*_CMY-2_

In contrast to the findings when examining the indicator *E. coli* collection from non-enriched samples, almost all ceftriaxone-enriched samples (n=145) harbored cephalosporin resistant *E. coli* (only one meconium sample was negative for the entire study) (Table 2). Almost all ceftriaxone-enriched samples demonstrating growth (n=144) harbored *E. coli* positive for *bla*_CMY-*2*_ except one sample after the ceftiofur cessation which only harbored *E. coli* positive for *bla*_CTX-M_. Although the *bla*_CTX-M_ gene was much less prevalent, it was present in all the production chain of both years. Few differences between flocks before and after the cessation of *in ovo* administration of ceftiofur were observed with respect to the proportion of samples harboring *E. coli* isolates positive for *bla*_CTX-M_, *bla*_TEM_ or *bla*_SHV_, namely the proportion of samples harboring *E. coli* positive for *bla*_CTX-M_ (p<0,05) or *bla*_TEM_ (p<0,01), which were increased for pooled feces of broilers for lincomycin-spectinomycin flocks compared to flocks with no *in ovo* antimicrobial administration. As observed for the non-enriched samples, the proportion of ceftriaxone-enriched samples harboring *E. coli* positive for *bla*_SHV_ was very low (results not show) and *bla*_OXA_ was not detected in any sample.

**Table 2:** Effect of cessation of *in ovo* administration of ceftiofur and the substitution with lincomycin-spectinomycin on the proportion of ceftriaxone-enriched samples from newly hatched, broiler and breeder birds with *E. coli* isolates positive for ESBL/AmpC resistance genes

| Sample origin | Before or after ceftiofur cessation in broiler | *In ovo* administration of: | No of samples with growth (%) after ceftriaxone enrichment | No of isolates<sup>a</sup> | % samples with one or more isolates positive for: |  |  |
| --- | --- | --- | --- | --- | --- | --- | --- |
|  |  |  |  |  | *bla*<sub>TEM</sub> | *bla*<sub>CMY-2</sub> | *bla*<sub>CTX-M</sub> |
| Meconium of newly hatched chicks | Before (2014) | Ceftiofur | 19 (95) | 57 | 26 | 100 | 16 |
|  | After (2015) | Nothing | 14 (100) | 40 | 43 | 93 | 7 |
|  | After (2015) | Lincomycin Spectinomycin | 16 (100) | 47 | 44 | 100 | 0 |
| Pooled feces of broilers | Before (2014) | Ceftiofur | 20 (100) | 60 | 40 | 100 | 20 |
|  | After (2015) | Nothing | 14 (100) | 42 | 7<sup>b</sup> | 100 | 0<sup>b</sup> |
|  | After (2015) | Lincomycin Spectinomycin | 16 (100) | 48 | 56<sup>c</sup> | 100 | 31<sup>c</sup> |
| Pooled feces of breeders | Before (2014) | Ceftiofur | 22 (100) | 66 | 45 | 100 | 14 |
|  | Before (2015) | Ceftiofur | 24 (100) | 72 | 67 | 100 | 8 |

<sup>a</sup> All isolates tested were uidA positive (likely *E. coli*). For each sample, 2-3 isolates were tested
<sup>bc</sup> Different letters in superscript in the same column and same sample indicate significantly different results

### 3. Antimicrobial non-susceptibility in ESBL/AmpC resistance gene positive *E. coli* isolates from ceftriaxone-enriched samples from newly hatched, broiler and breeder birds following the cessation of *in ovo* administration of ceftiofur and substitution with lincomycin-spectinomycin

Most (274/290) of the isolates from the ceftriaxone-enriched samples from all sources that were selected for antimicrobial susceptibility testing were positive for *bla*_CMY-2_ whereas 20/290 were positive for *bla*_CTX-M_. One isolate was negative for *bla*_CMY-2_ and *bla*_CTX-M_ but positive for *bla*_SHV_ and *bla*_TEM._ The isolates positive for *bla*_CTX-M_ were non-susceptible to a significantly lower number of different antimicrobials than those positive for *bla*_CMY-2_ (p<0,01) or both *bla*_CMY-2_ and *bla*_CTX-M_ (p<0,0001). Among the 15 different antimicrobials examined, the maximum number of antimicrobials for which non-susceptibility was observed in isolates positive for *bla*_CTX-M_ alone (n=15) was 8 whereas the maximum number of antimicrobials for which non-susceptibility was observed in isolates positive for *bla*_CMY-2_ alone (n=269) or for *bla*_CMY-2_ and *bla*_CTX-M_ (n=5) was 13 and 12 respectively.

Among samples from birds receiving the same antimicrobial regime *in ovo* at the hatchery, non-susceptibility for certain antimicrobials varied in the different production phases of the study. For flocks in which birds received ceftiofur *in ovo*, the proportion of samples with *E. coli* showing non-susceptibility in broilers was higher than in breeding flocks (p <0,001) and their meconiums (p <0,0001) for trimethoprim-sulphamethoxazole and than in breeding flocks (p = 0,02) for sulfisoxazole. No differences between phases were observed in flocks which received no antimicrobial *in ovo* following the cessation of *in ovo* administration of ceftiofur. However, for flocks which received lincomycin-spectinomycin *in ovo*, a greater proportion of non-susceptibility was observed in the broiler flocks compared to their breeders, for the aminoglycoside and folate inhibitor classes. For gentamicin (p <0,02) and spectinomycin (p <0,01), a greater proportion of non-susceptibility was found in meconiums compared to their breeders. For trimethoprim-sulphamethoxazole, a greater proportion of non-susceptibility was observed in broiler flocks compared to breeding samples (p <0,001). For sulfisoxazole, a greater proportion of non-susceptibility was observed in both meconium (p <0,01) and broilers (p <0.01) compared to breeding flocks.

### 4. Increased proportion of ceftriaxone-enriched samples with *E. coli* non-susceptible to antimicrobials of different classes following the cessation of *in ovo* administration of ceftiofur and substitution with lincomycin-spectinomycin

Thus, we observed that replacement of ceftiofur with lincomycin-spectinomycin for *in ovo* administration did not appear to affect the proportion of non-enriched samples harboring *E. coli* with the ESBL/AmpC resistance gene *bla*_CMY-2_. As enrichment with ceftriaxone showed that *E. coli* isolates with the ESBL/AmpC resistance gene *bla*_CMY-2_ are ubiquitous in most examined samples, we wished to determine the effect of lincomycin-spectinomycin administration on the non-susceptibility of these isolates to antimicrobials of the different categories and on the level of MDR. As expected, with almost 100% of isolates positive for *bla*_CMY-2_ gene, resistance to penicillins with and without β-lactamases inhibitors, cephalosporins of third generation, and cephamycin was almost 100%, irrespective of the antimicrobial used at the hatchery. There was no difference in the proportion of samples harboring *bla*_CMY-2_-positive *E. coli* non-susceptible for any of the six other antimicrobial classes in meconium or broiler feces between flocks before and after cessation of *in ovo* administration of ceftiofur when there was no replacement with lincomycin-spectinomycin, except for the non-susceptibility to trimethoprim-sulphamethoxazole in broiler feces where a significant decrease was observed following the ceftiofur cessation (p= 0,001).

On the other hand, there were a significantly greater proportion of samples harboring *bla*_CMY-2_-positive *E. coli* non-susceptible to streptomycin, trimethoprim-sulphamethoxazole, chloramphenicol, and tetracyclines in broiler feces when lincomycin-spectinomycin was administered *in ovo* than when no antimicrobial was used following cessation of administration of ceftiofur (Table 3). The same trend could be seen in the meconium, although the results were not statistically different, with the exception of spectinomycin. For many of these antimicrobials, a significantly greater proportion of samples harboring *bla*_CMY-2_-positive non-susceptible *E. coli* in meconium was observed when lincomycin-spectinomycin was administered *in ovo* than prior to cessation of administration of ceftiofur. A similar trend was observed in the broiler feces but was only significant for spectinomycin. Finally, neither the cessation of *in ovo* administration of ceftiofur nor replacement with lincomycin-spectinomycin affected the proportion of samples harboring *bla*_CMY-2_-positive *E. coli* non-susceptible to fluoroquinolones (nalidixic acid and ciprofloxacin) and macrolides (azithromycin). There was a very low proportion of non-susceptibility to these antimicrobials in isolates of the potential ESBL/AmpC-producing *E. coli* collection (Table 4).

**Table 3:**
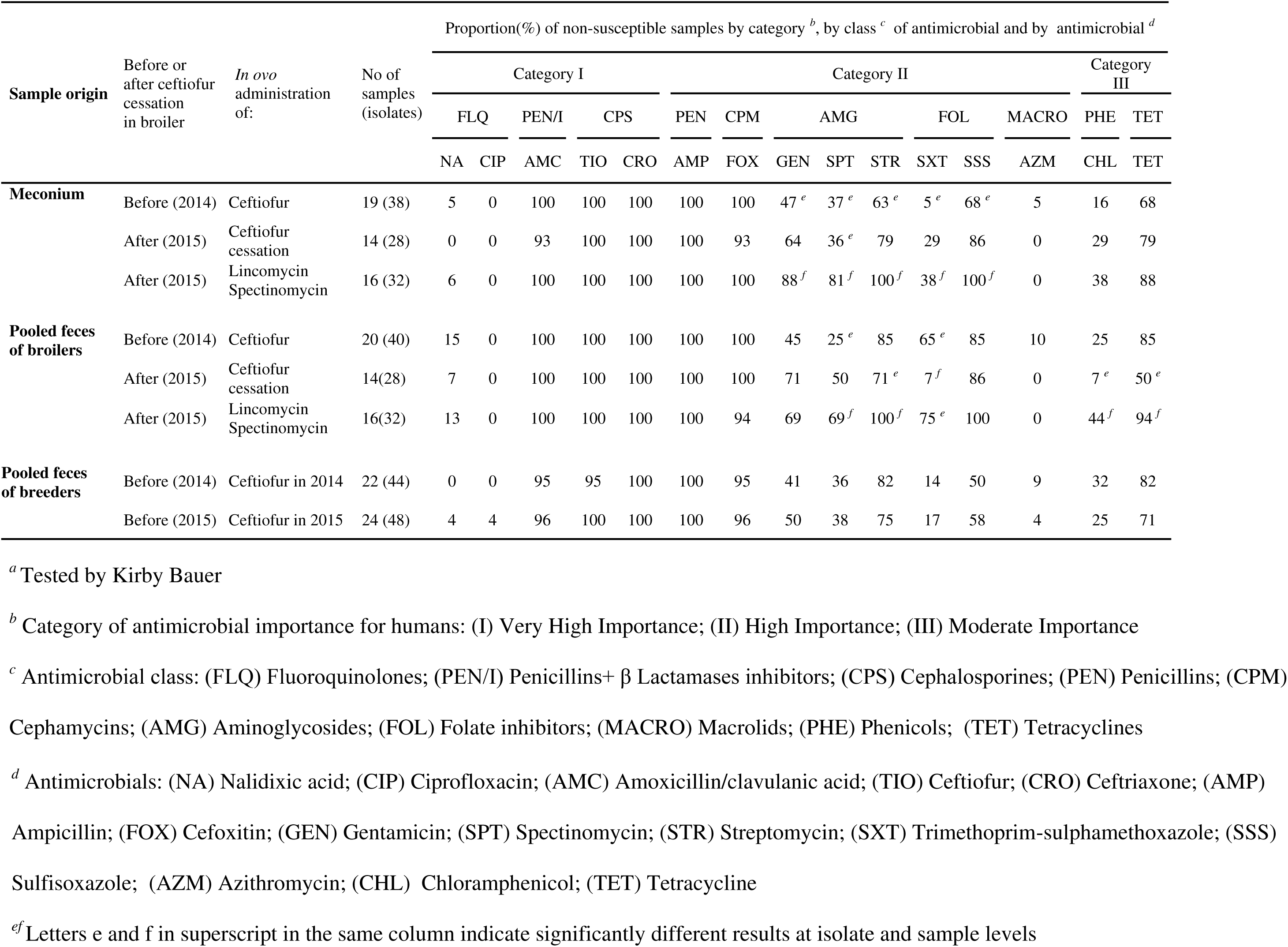
Effect of cessation of *in ovo* administration of ceftiofur and substitution with lincomycin-spectinomycin on the proportion of ceftriaxone-enriched samples from newly hatched, broiler and breeder birds harboring antimicrobial non-susceptible in ESBL/AmpC resistance gene positive *E. coli* samples *a*

**Table 4:**
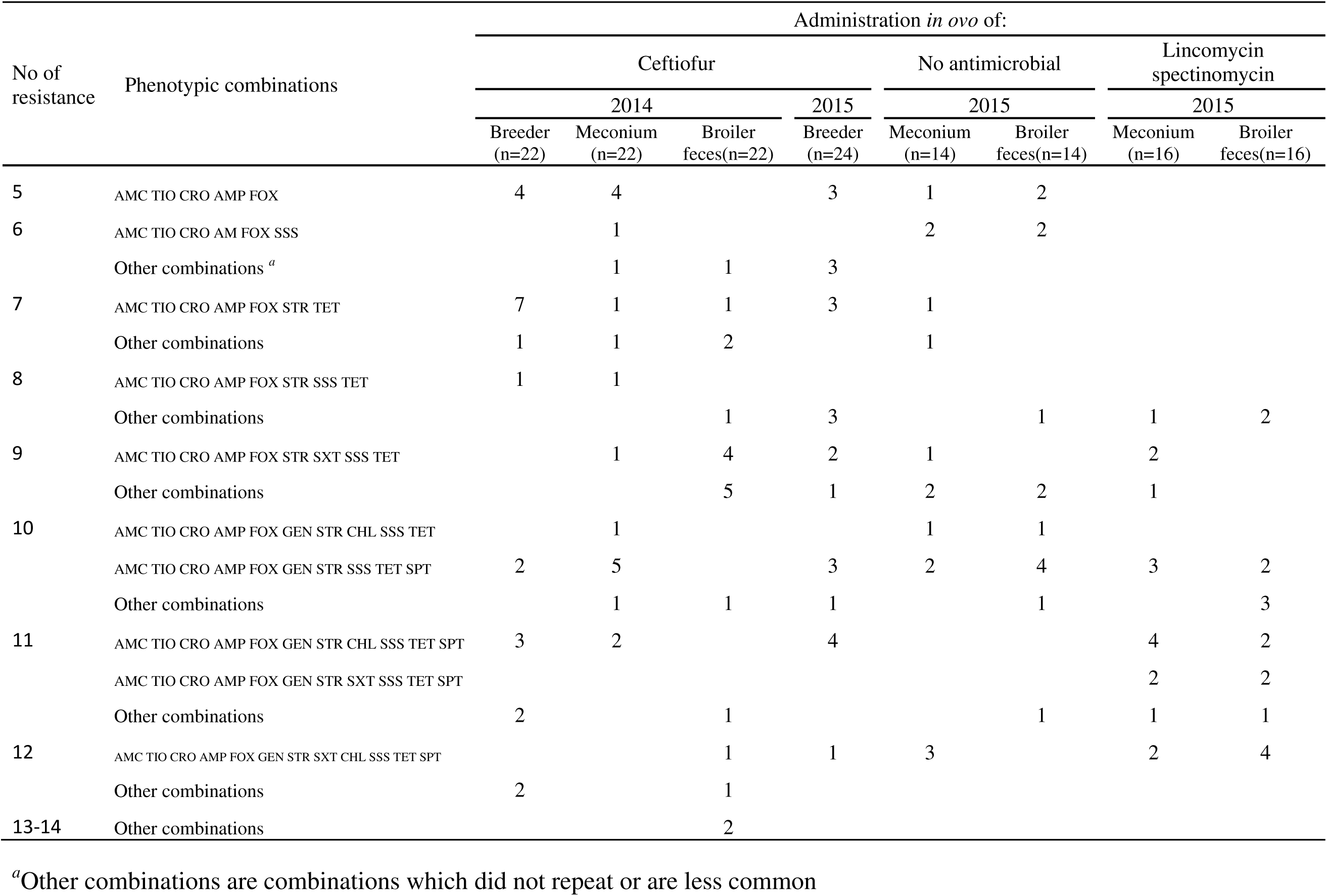
Effect of ceftiofur cessation and the substitution with lincomycin-spectinomycin on the proportion of antimicrobial phenotypic co-resistance profiles in samples from newly hatched, broiler and breeder birds with *E. coli* of the ceftriaxone enriched collection

### 5. Increased level of MDR in ESBL/AmpC resistance gene positive *E. coli* isolates in ceftriaxone-enriched samples from newly hatched, broiler and breeder birds following the cessation of *in ovo* administration of ceftiofur and substitution with lincomycin-spectinomycin

The proportion of ESBL/AmpC resistance gene positive *E. coli* isolates from ceftriaxone-enriched samples non-susceptible to 8 or more classes (referred to as possible extensively drug resistant (XDR) (27)) was greater in broiler feces than in the meconium in 2014 before the cessation of the *in ovo* administration of ceftiofur (Figure 1). The proportion of possible XDR isolates in breeder feces in 2014 before the cessation of ceftiofur was similar to that in broiler feces. The proportion of possible XDR isolates in broiler feces following cessation of ceftiofur when no antimicrobial was administered in ovo was slightly lower than that observed in broiler feces before the cessation of ceftiofur. In contrast, flocks receiving lincomycin-spectinomycin demonstrated a level of possible XDR similar to that that observed before ceftiofur cessation. Notably, all isolates in the lincomycin-spectinomycin treated flocks demonstrated non-β-lactam non-susceptibility in addition to β-lactam (penicillins with and without β-lactamases inhibitors, third generation cephalosporins and cephamycin) non-susceptibility (non-susceptibility being to 6 or more classes) whereas those in flocks prior to cessation of ceftiofur administration or following cessation of ceftiofur when no antimicrobial was administered *in ovo* often showed only β-lactam non-susceptibility (non-susceptibility being to 4 or more classes) (Table 4). Thus, samples from meconium and broiler feces of chicks having received lincomycin-spectinomycin *in ovo* showed non-susceptibility to a higher number of antimicrobial classes than those of chicks having received ceftiofur or which did not receive any antimicrobial *in ovo*. Finally, possible XDR bacteria in flocks that had received lincomycin-spectinomycin almost always demonstrated non-susceptibility to spectinomycin and were mostly associated with non-susceptibility to other non-β-lactams such as gentamicin, streptomycin, sulfisoxazole and tetracycline.

**Figure 1:**
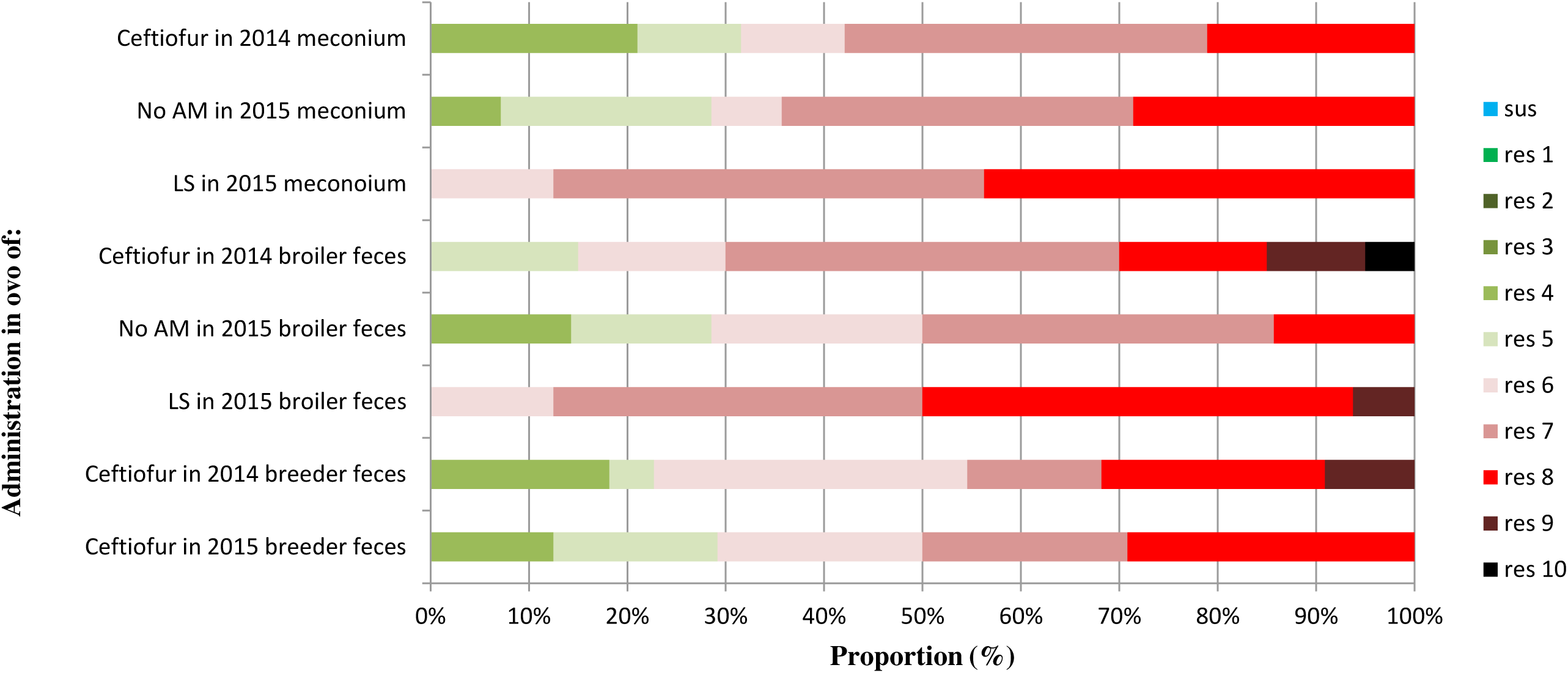
Effect of cessation of *in ovo* administration of ceftiofur and replacement with lincomycin-spectinomycin on the proportion of multidrug resistance in ESBL/AmpC resistance gene positive *E. coli* isolates in ceftriaxone-enriched samples from newly hatched, broiler and breeder birds.

LS: Lincomycin spectinomycin; No AM: no antimicrobial *in* ovo; sus: susceptible to all antimicrobial classes; res1-res10: resistance to one up to ten different antimicrobial classes, 2014 is before and 2015 is after the cessation of ceftiofur in Canada

## Discussion

Examination of the indicator *E. coli* in our study clearly demonstrates a decrease in prevalence of samples harboring isolates positive for the ESBL/AmpC resistance genes *bla*_CMY-2_ and *bla*_CTX-M_ in the meconium of newly hatched birds after the cessation of in ovo administration of ceftiofur. This finding shows a beneficial effect of the cessation of the use of ceftiofur, reinforcing the findings of other studies where a decrease in resistance to extended spectrum cephalosporins was observed (4, 9, 12). Nevertheless, we did not observe an effect in the broilers, as was the case in the previous studies (7). In addition, the substitution of ceftiofur with lincomycin-spectinomycin in the broiler hatchery did not have a significant impact on the presence of *bla*_CMY-2_ or *bla*_CTX-M_ in this collection, reinforcing the idea that these genes were selected mainly due to the use of ceftiofur. When we used a more sensitive approach of enrichment with ceftriaxone, we found that almost all samples, irrespective of the production phase, harbored isolates positive for the ESBL/AmpC resistance genes, mostly *bla*_CMY-2,_ before the cessation of the use of ceftiofur, and this was not affected by the cessation, at least in the first year after it occurred.

We observed *E. coli* positive for *bla*_CTX-M_ across all the poultry production chain, suggesting that this gene is more prevalent in Canadian flocks than previously reported (6). Our results following ceftriaxone enrichment demonstrated that cephalosporin resistance genes were still present among the population but that the prevalence was probably lower in the flocks following the cessation of use of ceftiofur. The lower prevalence of *bla*_CTX-M_ with respect to *bla*_CMY-*2*_ in our study in contrast to the lower prevalence of *bla*_CMY-*2*_ reported in Europe (5) could be due to the presence of different plasmids containing *bla*_CTX-M_ in Europe such as IncN, IncI, IncL/M and IncK (8). Nevertheless, a more recent study in Europe showed a higher proportion of *bla*_CMY-2_ (28). The difference in relative predominance of the *bla*_CTX-M_ and *bla*_CMY-2_.genes could also be explained in part by differences in geographical distribution of these genes (7).

Although the *in ovo* administration of ceftiofur ceased in Canada in 2014, this practice was only stopped in parent flocks in the US in 2015 after the present study had been completed; hence, vertical transmission of resistance genes may have occurred (7, 29, 30). In fact, the results of our study are similar to those obtained in Denmark, where the parent flocks which did not receive ceftiofur but for which the grandparent flocks received cephalosporins (19), demonstrated a 93% prevalence of cephalosporin resistant *E. coli* (most of the samples being positive for *bla*_CMY-2_). Vertical transmission of APEC from breeders to their progeny is well known (2) and has been demonstrated recently in the Nordic countries where APEC ST117 O78:H4 was transmitted from grand-parent flocks to their broilers in different countries (31). In addition, Nilsson et al. demonstrated the vertical transmission of *E. coli* carrying *bla*_CMY-2_ from grand-parent flocks which had been exposed to ceftiofur to parent and broiler flocks which had never received ceftiofur (29). Zurfluh et al. demonstrated similar results where *E. coli* carrying *bla*_CTX-M_ were vertically transmitted from parent flocks to hatchery and broiler flocks (30). Horizontal transmission has also been reported for ESBL/AmpC-associated resistance genes which are found on horizontally transferable plasmids (32). Thus, the administration of ceftiofur *in ovo* to US breeders could have resulted in an increase in the proportion of resistance to third generation cephalosporins in broiler *E. coli* even though the administration of this antimicrobial had ceased in Canadian hatcheries. In the Danish study, a decrease in prevalence of samples positive for cephalosporin resistant *E. coli* to 27% in broiler flocks originating from breeders which did not receive ceftiofur (19). Finally, maintenance of the high proportion positive samples in the Canadian broilers at one year after the cessation of administration of ceftiofur could also be explained by environmental contamination which may have persisted after birds ceased to excrete *E. coli* positive for ESBL/AmpC resistance genes (28).

Our finding of an increased prevalence of non-susceptibility of *E. coli* to the aminoglycosides, phenicols, the tetracyclines and the folate inhibitors in 2015 for the meconium and broiler feces of birds receiving lincomycin-spectinomycin clearly showed that the use of this combination resulted in an increase in MDR among the cephalosporin resistant *E. coli*. All samples from birds receiving lincomycin-spectinomycin harbored isolates with non-susceptibility to at least 6 classes whereas those which had received ceftiofur before its cessation or no antimicrobial *in ovo* harbored isolates non-susceptible to only 4 classes, the gene *bla*_CMY-2_ conferring non-susceptibility to the 4 classes of β-lactams. These results suggested that there may be a co-selection phenomenon when using lincomycin-spectinomycin, which would explain the high MDR for this group. The association of non-susceptibility to spectinomycin with non-susceptibility to other non-β-lactams such as gentamicin, streptomycin, trimethoprim-sulphamethoxazole, sulfisoxazole, chloramphenicol and tetracyclines could be due to the presence of a plasmid or transferable elements (integrons, transposons, insertion sequences) that carry *bla*_CMY-2_ and other resistance genes in addition to those encoding resistance to spectinomycin. In a study of *E. coli* isolates from clinical cases of colibacillosis in chickens in Québec, Canada, Chalmers et al (6) demonstrated that the use of spectinomycin could increase co-selection. Indeed, the presence of resistance to spectinomycin (*aac(3)-VI* gene) and gentamicin (*aadA)*, an antimicrobial that is no longer routinely used in Canadian poultry, were strongly associated statistically and located on a modified class 1 integron. These authors also demonstrated that the use of spectinomycin did not seem to select cephalosporin resistant *E. coli* or APEC. The *aac(3)-VI* and *bla*_CMY-2_ genes were negatively associated, but when present together, were generally located on the same plasmids (6). An increase of bacteria with co-selection of antimicrobial resistance and greater MDR would thus be expected with the use of lincomycin-spectinomycin and could allow the emergence of a bacterial flora that contains more MDR and, possibly, XDR bacteria.

In a study of MDR *E. coli* harboring *bla*_CMY-2_ or *bla*_CTX-M_ from sick pigs in Canada, Jahanbakhsh et al (11) demonstrated the presence of class 1 integrons which were significantly associated with resistance to gentamicin, kanamycin, streptomycin, trimethoprim/sulfamethoxazole, sulfisoxazole, chloramphenicol and tetracycline. This study also demonstrated the presence of an IncA/C plasmid positively associated with resistance to gentamicin, kanamycin, streptomycin, trimethoprim-sulfamethoxazole, sulfisoxazole, chloramphenicol and tetracycline, in *E. coli* with resistance to at least one antimicrobial of eight different classes, the presence of *bla*_TEM_ and class 1 integron. Furthermore, other plasmids such as IncFIB had a positive association with IncA/C whereas IncI1 plasmid had a negative association with these plasmids (11). The presence of such plasmids in our study may explain why multidrug-resistant bacteria were selected when using lincomycin-spectinomycin at the hatchery. Studies in Japan and Netherlands also found IncA/C plasmids harboring *bla*_CMY-2_ and resistance to non β-lactam antimicrobial (28, 33). The plasmids selected in our study may have contained *bla*_CMY-2_ genes and other resistance genes. Alternatively, certain clonal lineages may harbor one plasmid with the *bla*_CMY-2_ genes concomitantly with a second plasmid conferring resistance to non β-lactam antimicrobials. Plasmids of several incompatibility groups, such as as IncIγ, IncB/O, IncFIB, IncFIC and IncI1, have already been reported to contain *bla*_CMY-2_ genes alone, but not resistance to non β-lactam antimicrobials (11, 33).

Horizontal and vertical transmission of *bla*_CMY-2_ has already been reported (19), thus further testing is needed to evaluate if vertical transmission of *bla*_CMY-2_ and *bla*_CTX-M_ occurs between the breeders and broilers of our study.

We observed a significant decrease in non-susceptibility to trimethoprim-sulphamethoxazole in the broiler feces for the flocks which did not receive any antimicrobial *in ovo* following the cessation of ceftiofur, thus demonstrating the potential for a positive cessation effect for this class of antimicrobials. This decrease could be explained by a selective disadvantage of *bla*_CMY-2_ on certain plasmids at the animal level when ceftiofur is stopped and no antimicrobial is used subsequently (28). Furthermore, resistance to phenicols and tetracyclines were lower, although not significantly, for these same flocks, showing a beneficial effect with respect to the prevalence of MDR. Nevertheless, no significant differences were seen for the meconium and broiler feces between flocks receiving ceftiofur prior to its cessation and flocks when ceftiofur is stopped and no antimicrobial is used subsequently with respect to non-susceptibility to gentamicin, spectinomycin, sulfisoxazole, tetracyclines and chloramphenicol, which suggests that resistance to these antimicrobials is mainly due to the use of lincomycin-spectinomycin. However, antimicrobials received during rearing may also affect the variation in the prevalence of non-susceptibility to these antimicrobials. If broiler flocks not receiving antimicrobials *in ovo* have lower level of resistance to certain antimicrobials (such as trimethoprim-sulphamethoxazole), stopping antimicrobials at the hatchery could lead to clinically more effective antimicrobial treatments during fattening.

We observed very little non-susceptibility to the fluoroquinolones (nalidixic acid and ciprofloxacin) and macrolids (azithromycin) in *E. coli* isolates obtained following ceftriaxone enrichment. This result was expected as these classes of antimicrobial agent were not approved for use in poultry in Canada during the years of our study, during which time ceftiofur, bacitracin, virginiamycin, avilamycin, and lincomycin-spectinomycin were the most commonly used antimicrobials(3). In addition, ceftriaxone enrichment is not a selective method for detection of quinolones or macrolids resistance. Finally, resistance to quinolones is mainly due to the acquisition of chromosomal mutations rather than to resistance genes present on plasmids as is the case for ESBL/AmpC production (34).

Overall, our results demonstrate a decrease in the prevalence of cephalosporin-resistant *E. coli* harboring *bla*_CMY-2_ and *bla*_CTX-M_ after the ceftiofur cessation and replacement with lincomycin-spectinomycin. No differences were seen in the proportion of *bla*_SHV_, *bla*_OXA_ and *bla*_TEM_. On other hand, when samples were examined using the more sensitive ceftriaxone-enrichment, 99% of samples were shown to harbor cephalosporin-resistant *E. coli*, indicating that at the sample level, there has not yet been a reduction in the prevalence of *E. coli* carrying the ESBL/AmpC resistance genes one year after the ceftiofur cessation. The 2015 ceftiofur cessation in USA breeders could possibly decrease this proportion in the future. Also, we observed a high proportion of multidrug resistant ESBL/AmpC *E. coli* in young chicks, broilers and breeders, with an increase in the proportion of *E. coli* possibly extensively resistant in flocks receiving lincomycin-spectinomycin, which is a concern for public health.

## Acknowledgments

This work was supported by Agriculture and Agri-Food Canada and by Éleveurs de volailles du Québec, Agri-Innovation Program grant AIP P270. We thank Ghyslaine Vanier, Gabriel Desmarais, and Jocelyn Bernier-Lachance for their technical assistance, Benoît Lanthier for assistance in the field and Les Fonds du Centenaire of the faculty of veterinary medicine of Université de Montréal for their contribution.

